# HIF1α and HIF2α mRNA Expression in Oral Squamous Cell Carcinoma does not correlate with protein expression

**DOI:** 10.1101/2025.05.02.651922

**Authors:** Pooja Singh, Manoj Pandey, Gopal Nath, Deepak Kumar, Amrita Ghosh Kar

**Author notes:** **Address for correspondence-** Prof. Manoj Pandey, Surgical Oncology, Banaras Hindu University Institute of Medical Science, Varanasi, India, 221005.

## Abstract

**Background:** Oral cancer is one of the common cancers among men, its presentation is usually late. Tumor growth leads to hypoxia, and HIF1α and HIF2α are important regulators of hypoxia induced pathways, inducing angiogenesis and tumor growth. This study compares the expression levels of HIF1α and HIF2α mRNA in OSCC tissues to adjacent normal mucosa.

**Methods:** RNA was isolated from 85 OSCC tissues and adjacent normal mucosa followed by real-time (RT) PCR to quantify HIF1α and HIF2α expression using GAPDH and β-actin respectively as reference genes.

**Results:** HIF1α overexpression was detected in 76.47% of case and 71.76% of controls (p=0.4906), whereas HIF2α overexpression occurred in 91.76% of cases 94.11% of controls (p=0.35). HIF1α and HIF2α expression level were not significantly different in OSCC tissues compared to adjacent normal mucosa.

**Conclusion:** HIF1α and HIF2α play important role in hypoxia induced pathways and tumor progression, yet this study found no significant variation in expression between OSCC tissues and surrounding normal tissues. This suggest that their expression may be due to physiological adaptability to hypoxia or stress, or may be due to use of adjacent tissue as control and loss of protein expression could be due to post translational modifications.

## Introduction

Cancers of the oral cavity are a substantial global health concern, with 389,485 new diagnoses and 188,230 fatalities in 2022 (1), this number is expected to rise by 10-12% by 2025. Hypoxia has been implicated in oral cancer and various mechanisms have been proposed (2, 3). The transcription factors HIF1α and JMJD1A have been validated as key hypoxia dependent regulators in the carcinogenesis of HNSCC (4), with the HIF1α and HIF2α identified as overexpressed biomarkers in various other cancers, like papillary thyroid carcinoma (5). The regulatory connection between HIF2α and VEGFR-1, VEGFR2 and MMP2 through the ETS1/HIF2α has also been identified (6). However, in HPV positive oropharyngeal tumors elevated VEGF mRNA level have been described through a HIF-1 independent mechanism (7). Gene alterations may regulates HIF1α expression under nonhypoxic conditions in HNSCC, potentially priming cells for hypoxia tumor environments (8). Although angiogenic markers such as miR-210 and HIF1α are unreliable indicator of malignancy in salivary glands, LMP1(latent membrane protein 1) directly upregulates HIF1α transcription and post transcription in nasopharyngeal carcinoma (NPC) cells, increasing HIF1α mRNA level even under normoxic condition (9). Inhibiting HIF1α expression reduces the aggressive potential of SCC-15 cell lines under both normoxic and hypoxic conditions, by inhibiting angiogenesis, glycolysis and metastasis, making it a potential strategy to improve survival in tongue squamous cell carcinoma (TSCC) (10). Furthermore, EGFR inhibitors like gefitinib and erlotinib reduce VEGF expression via distinct mechanism including reduced Sp1 binding to the VEGF promoters and HIF1α downregulation (11), while TSA (trichostatin) exhibit anticancer effects in tongue SCC cells by inhibiting HIF1α and VEGF expression (12). A biochemical link between tumor hypoxia and neoangiogenesis has been demonstrated through hypoxia induced regulation of VEGF and HIF1α expression (13), with HIF1α is shown to suppress proliferation and apoptosis under hypoxia condition in Tca8113 cells via activation of the Hippo signaling pathway (14) (Figure 1) The hippo pathway primarily involved a cascade from Hippo to oncoprotein YAP / TAZ kinase (15). It has been found TAZ levels were strongly linked to tumor occurrence, development and prognosis (16, 17). Elevated nuclear prolyl hydroxylase (PHD2) level are associated with poorly differentiated and highly proliferative tumors, indicating insufficient HIF1α downregulation (18). Angiogenin, upregulated in the hypoxic microenvironment of oral cancers, holds therapeutic potential through its inhibition,(19) while resveratrol significantly inhibits basal and hypoxia induced HIF1α protein accumulation in cancer cells without affecting mRNA level (20). VEGF-A expression, frequently upregulated in human malignancies and associated with hypoxia markers, can also increase independent of hypoxia, highlighting the complexity of hypoxia related pathways in cancer progression (21). Majority of the studies on hypoxia are in cell lines, with little or no human studies. This study was carried out to measure the mRNA of HIF1α and HIF 2α expression in oral cancer and normal tissue.

**Figure 1:**
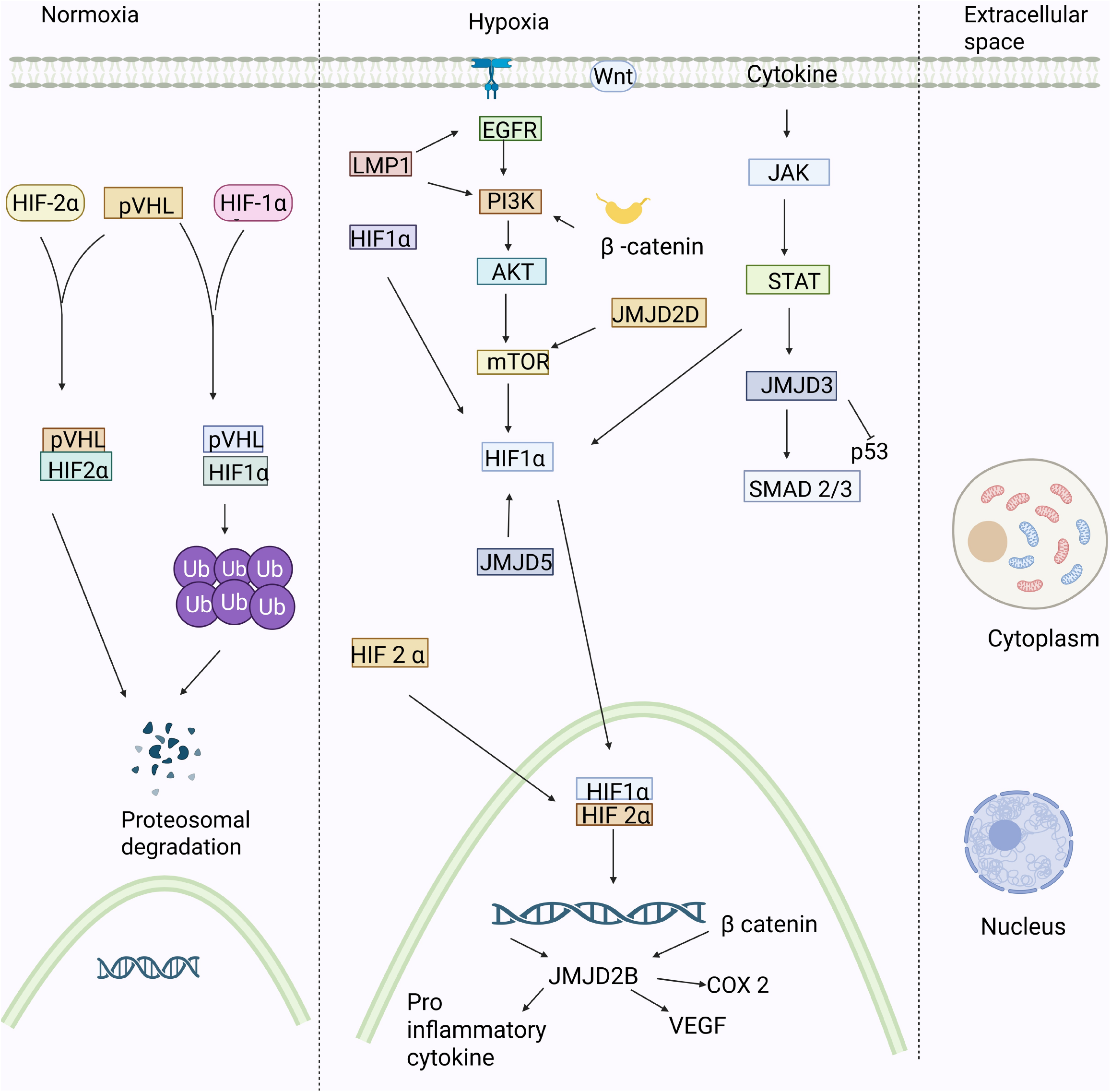
Showing degradation of HIF under normaxia and HIF downstream pathway under hypoxia

## Material and methods

### Patient selection

Oral cancer tissue samples were collected from histologically confirmed 85 patients diagnosed with oral squamous cell carcinoma after obtaining written informed consent and approval by the ethics committee. Specimens were obtained from the primary tumor sites during surgical resection, ensuring representation of the pathological characteristic of the cancerous tissue. Additionally, normal oral mucosa sample were collected from adjacent site after tumor resection as control for comparison

### RNA Extraction Procedure

RNA extraction was carried out using the Pure Link™ RNA Mini Kit Invitrogen™ RNA Extraction kit after homogenizing a 10 mm of tissue sample using a motor and pestle. After through mixing 5 ml of 1XPBS was added, and the sample was transferred to a 15 ml tube. After Centrifuging at 500 X g (4500 rpm) for 10 minutes at 4°C the pellet was separated, and the supernatant was discarded. Complete lysis was achieved by adding 600 μl of lysis buffer and mercaptoethanol to the sample and mixing thoroughly by vortexing. After adding an equal volume of 70% (600 μl) ethanol to the lysate and mixing thoroughly 700 μl of the lysate ethanol mixture was transferred to the spin cartridge in a collection tube and centrifuge at 12,000 rpm for 2 minutes. Sample was washed with 700 μl wash buffer to the cartridge, centrifuged at 12,000 rpm for 1 minute. For elution, the sample was transferred from the spin cartridge to a clean RNase free collection tube, and 40μl of RNase free water was added directly to the membrane, incubate for 1 minute at room temperature and centrifuge at 12,000 rpm for 2 minute to elute the RNA. Eluted RNA was stored at −20 °C

### Quantification of isolated total RNA

RNA samples were quantify using a spectrophotometer (Thermo Fisher) at 260 nm and purity of RNA was checked by ratio of 260/280 nm and RNA samples having ratio 1.8-2.0 were checked for qualitative analysis to determine their concentrations in nanograms per microliter (ng/μl). The concentration was then normalized to ensure uniform RNA input for subsequent experiments. To normalize, the concentration of each sample was adjusted to using nuclease free water. The amount of RNA required for normalization was calculated using the formula V1 = C2 X V2/C1.

### Real-Time PCR (RT-PCR)

For each reaction, 6.8 μl of nuclease free water was added followed by 10 μl of Syber Green Master Mix (Brilliant III Ultra-Fast SYBR® Green QRT-PCR Master Mix), then 1μl of forward and reverse primer was added (Eurofins). The primer sequences used were HIF-1α - Forward CCAGTTACGTTCCTTCGATCAG Reverse GTAGTGGTGGCATTAGCAGTAG, GAPDH-Forward CAAGAGCACAAGAGGAAGAGAG Reverse CTACATGGCAACTGTGAGGAG, HIF2α-Forward GGCTGTGTCTGAGAAGAGTAAC Reverse CCCGAAATCCAGAGAGATGATG, Beta-actin-Forward CACTCTTCCAGCCTTCCTTC Reverse GTACAGGTCTTTGCGGATGT.

This was followed by 0.2 μl of DTT (dithiothreitol) to stabilize the enzyme, followed by 1 μl of reverse transcriptase (RT) for converting RNA into complementary DNA (cDNA). Finally, 1 μl of normalized RNA sample was added to the mixture. The PCR plate was sealed with optical caps to prevent evaporation and centrifuged to ensure that all reagents are collected at the bottom of the wells.

The reaction is prepared load them into (Bio-Rad CFX96) real time PCR machine. The cycler was programed to include one reverse transcription steps at 50 °C for 10 min, allowing the RNA to be reverse transcribed into cDNA. This was followed with denaturation step at 95° C for 3 min to denature the cDNA strands. The amplification stage consists of 40 cycles, with each cycle including a denaturation step at 95° C for 5 sec and an annealing/ extension step at 60° C for 10 sec.

After completing the run the Ct (threshold cycle) values were analyzed and the melting curve was examined to confirm the specificity of the amplification. The data is shown as numbers and percentages calculated from the ratio of house keeper gene and the HIF 1 and 2 gene amplification. The Chi-square test was used for statistical analysis.

## Results

Of the 85 patients studied, there were 76 male (89%) and 9 females (10%). There was no case in stage I, while 8 cases (9.4%) were in stage II. Majority of the patients were locally advanced being in stage III in 14 cases (16.4%) and stage IVA has the highest in 27 cases (31%) and IVB has the 2 cases (2.35%) (Table 1). Majority of tumors were located in the buccal mucosa (BM) (28%), 11% in the tongue and 20% in the alveolus. Tumor grades showed that 17.6% was well differentiated (WD), 31% moderately differentiated (MD)and 1.17%, poorly differentiated (PD) with no undifferentiated (UD).

**Table 1:** Patient characteristics for Cases and Adjacent Normal Tissue.

|  |  | Case and adjacent normal |  |
| --- | --- | --- | --- |
|  |  | n | % |
| Age (mean ± SD) |  |  |  |
| Gender | Male | 76 | 89 |
|  | Female | 9 | 10 |
| **Stage** | **I** | **0** | **0** |
|  | **II** | **8** | **9.4** |
|  | **III** | **14** | **16.4** |
|  | **IVA** | **27** | **31.76** |
|  | **IVB** | **2** | **2.35** |
| **Site** | **BM** | **24** | **28** |
|  | **Tongue** | **10** | **11** |
|  | **Alveolus** | **17** | **20** |
|  | **Other** |  |  |
| **Grade** | **WD** | **15** | **17.6** |
|  | **MD** | **27** | **31** |
|  | **PD** | **1** | **1.17** |
|  | **UD** |  |  |

Overexpression of HIF-1α was found in 76.4% of cases (65 individuals) and 71.7% of controls (61 individuals), while under expression was found in (23.5%) of 20 cases and 28.2% of controls (24). There was no statistically significant difference in HIF1α expression levels between patients and controls (p=0.48). Similarly, HIF2α overexpression occurred in 91.7% of cases (78 individuals) and 94.1% of controls (80 individuals), while under expression was detected in 7 cases (8.2%) and 5 controls (5.8%) (p-0.54). The table 2 compares the expression levels of two hypoxia inducible factors, HIF1α and HIF2α, between case and control groups, showing both overexpression and under expression rates.

**Table 2:** Comparison of HIF1 α and HIF2α Expression Levels Between Case and Control Groups.

| **Sample Type** |  | **Case**<br>**Over expressed** |  | **Control** |  | **p-value** |
| --- | --- | --- | --- | --- | --- | --- |
|  |  | **N** | **%** | **N** | **%** |  |
| HIF-1α | Over | 65 | 76.47 | 61 | 71.76 | 0..48 |
|  | Under | 20 | 23.52 | 24 | 28.23 |  |
| HIF-2α | Over | 78 | 91.76 | 80 | 94.11 | 0..54 |
|  | Under | 7 | 8.23 | 5 | 5.88 |  |

The table 3 shows data on HIF1 and HIF2 overexpression in various groups according on gender, tumor stage, tumor site and tumor grade. No statistically significant difference was observed either in cases or control for both HIF 1 and HIF 2α.

**Table 3:** Distribution of HIF1 and HIF2 Overexpression and Underexpression across different clinical and demographic variables.

| Variable |  | HIF 1 control |  | P<br>value | HIF 1<br>Case |  | P<br>value | HIF 2<br>control |  | P<br>value | HIF 2 cases |  | P<br>value |
| --- | --- | --- | --- | --- | --- | --- | --- | --- | --- | --- | --- | --- | --- |
|  |  | Over | Under |  | Over | Under |  | Over | Under |  | Over | under |  |
| Gender | Male | 52 | 21 | 0.4 | 53 | 18 | 0.18 | 69 | 2 | .1 | 64 | 2 | .1 |
|  | Female | 8 | 1 |  | 0 | 1 |  | 0 | 9 |  | 0 | 9 |  |
| Stage | II | 1 | 3 | .07 | 2 | 3 | .07 | 4 | 0 | .79 | 4 | 0 | .68 |
|  | III | 5 | 0 |  | 5 | 0 |  | 5 | 0 |  | 5 | 0 |  |
|  | Iva+b | 20+7 | 5+7 |  | 7+21 | 4+4 |  | 10+25 | 1+1 |  | 7+23 | 1+1 |  |
| Site | BM | 14 | 9 | .39 | 14 | 7 | .37 | 21 | 1 | .73 | 19 | 1 | .68 |
|  | Tongue | 6 | 3 |  | 7 | 1 |  | 9 | 0 |  | 9 | 0 |  |
|  | Alveolus | 13 | 3 |  | 14 | 3 |  | 14 | 1 |  | 11 | 1 |  |
|  | Others |  |  |  |  |  |  |  |  |  |  |  |  |
| Grade | Well | 9 | 6 | .49 | 9 | 3 | .82 | 13 | 1 | .88 | 12 | 1 | .89 |

The table 4 compares the immunohistochemistry (IHC) expression of HIF1α and HIF2α proteins and their related mRNA levels in control and case samples. Among the controls with HIF1α immunoexpression four cases showed mRNA overexpression and five showed underexpression with a p value of 0.05 indicating a statistically significant difference. Majority of the patients where protein expression was absent (56) showed mRNA overexpression implying that HIF1α protein is not detectable using IHC due to the post transcriptional regulation or protein breakdown. Only 3 cases showed both mRNA and protein expression.

**Table 4:** Comparison of Mrna Expression and IHC protein Expression of HIF1α and HIF2α in Control and Case sample.

| IHC<br>expression | MRNA→ | HIF 1 control |  | P value | HIF 1 Case |  | P<br>value | HIF 2 control |  | P value | HIF 2 cases |  | P<br>value |
| --- | --- | --- | --- | --- | --- | --- | --- | --- | --- | --- | --- | --- | --- |
|  |  | Over | Under |  | Over | Under |  | Over | Under |  | Over | under |  |
| HIF 1Con | Present | 4 | 5 | 0.05 | 3 | 5 | .005 | 7 | 0 | 1.0 | 7 | 0 | .64 |
|  | Absent | 56 | 17 |  | 58 | 13 |  | 66 | 7 |  | 66 | 2 |  |
| HIF 1 Case | Present | 3 | 3 | .18 | 2 | 3 | .04 | 8 | 0 | .63 | 5 | 0 | 1.0 |
|  | Absent | 57 | 19 |  | 59 | 15 |  | 70 | 2 |  | 68 | 2 |  |
| HIF2 con | Present | 4 | 5 | 0.05 | 3 | 5 | .005 | 8 | 0 | .63 | 7 | 0 | .65 |
|  | Absent | 56 | 17 |  | 58 | 13 |  | 70 | 2 |  | 71 | 2 |  |
| HIF 2 case | Present | 4 | 4 | .11 | 3 | 4 | .023 | 2 | 0 | 1.0 | 7 | 0 | 1.0 |
|  | Absent | 56 | 18 |  | 58 | 14 |  | 66 | 2 |  | 66 | 2 |  |

In HIF2α control samples 7 samples showed both mRNA and protein overexpression, in the rest of the cases mRNA was present but protein expression was not identified.

## Discussion

This study examines the mRNA expression levels of hypoxia inducible factors (HIFs) in oral squamous cell carcinoma (OSCC) tissues against normal adjacent mucosa. Even though HIF1α and HIF2α play important role in cancer progression, angiogenesis and hypoxia induced pathways, there were no significant variations in their expression levels between the case and control groups both showing over 70% expression for HIF1α mRNA and over 90% expression of HIF 2α. These results suggest that over expression of mRNA for both HIF1α and HIF2α expression is not limited to malignant tissues, but reflects physiological adaption or stress in surrounding normal tissues, as well. HIF1α overexpression in OSCC is linked to increased tumor size, advanced stage, lymph node metastasis and decreased overall survival making it a viable independent prognostic marker (22), however in the present study no relationship was observed with pathological variables.

In other tumors, higher expression has been associated with metastasis in prostate cancer cell lines and in renal carcinoma (23). In gastric cancer, targeting HIF1α, GLUT1 and LDH-5 was found to play essential roles in tumor energy metabolism (24) and was found to be associated with lymph node metastases, tumor infiltration depth, advanced TNM stages, and poor prognosis (25). Previous research has shown that HIF1α and HIF2α promote angiogenesis and tumor growth via regulating VEGF and modulating hypoxia related pathways. However, our data suggests that gene mutation, epigenetic changes, or normoxic HIF regulation, may contribute to OSCC formation.

HNSCC gene changes can influence HIF1α expression in non-hypoxic condition potentially preparing cells for future hypoxic situations. The function of LMP1 in nasopharyngeal cancer and potential to upregulate HIF1α under normoxia highlights the need to investigate alternate HIF regulatory routes. Similarly, HIF2α expression at mRNA and protein levels correlated with vascular and inflammatory histomorphological features in clear cell renal cell carcinoma (ccRCC) patients (26), offering prognostic value and insights into targeted therapy response. In ovarian cancer cell lines, immunohistochemistry showed elevated HIF1α and MMP13 in both tumor and metastasis with hypoxia increasing their mRNA and protein levels. SiRNA silencing of HIF1α reduced MMP13 expression and A2780 cell invasion, highlighting the role of hypoxia in driving cancer progression (27). Cells adapt to hypoxia by shifting from general function to specialized hypoxia response pathways with HIFs regulating both transcription and translation. Hypoxia slows canonical protein synthesis due to limited ATP, necessitating alternative mechanisms that contribute to cancer hallmarks like treatment resistance (28). HIF1α and its target gene, VEGF were up regulated in intraperitoneal tumors but not in primary gastric malignancies. Different microenvironment may alter the expression of these genes (29).

High mRNA levels and low protein expression in HIF1α and HIF2α in oral cancer as has been seen in this study can be explained by numerous molecular mechanisms including mRNA mutation, miRNA, post translational modification and protein stability factors. Post translational modification (PTMs) regulates the stability and function of HIF1α allowing for regulated expression under different cellular circumstances. Under normoxic circumstances, HIF1α is rapidly degraded due to hydroxylation of particular proline residues (Pro402 and Pro564) (24, 30–32). This alteration unable recognition by the von Hippel Lindau (VHL) proteins, resulting in ubiquition mediated proteosomal destruction (Figure 2). Ubiquitination is a major degradation signal that marks HIF1α for destruction in proteosome preventing buildup even with high mRNA expression levels (33, 34). Additionally, SUMOylation and acetylation have a substantial impact on protein stability and function. SUMOylation can stabilize or degrade HIF1α while acetylation affects its transcriptional activity and interaction with other proteins (35). These PTMs HIF1α availability preventing excessive activity in normoxia and stabilizing it under hypoxic condition, ensuring cellular homeostasis methylation and histone alteration serve critical role in regulating gene expression at both the transcriptional and translation levels (36-38). Promoter methylation which is usually associated with gene silence can sometime enhanced mRNA transcription.

**Figure 2:**
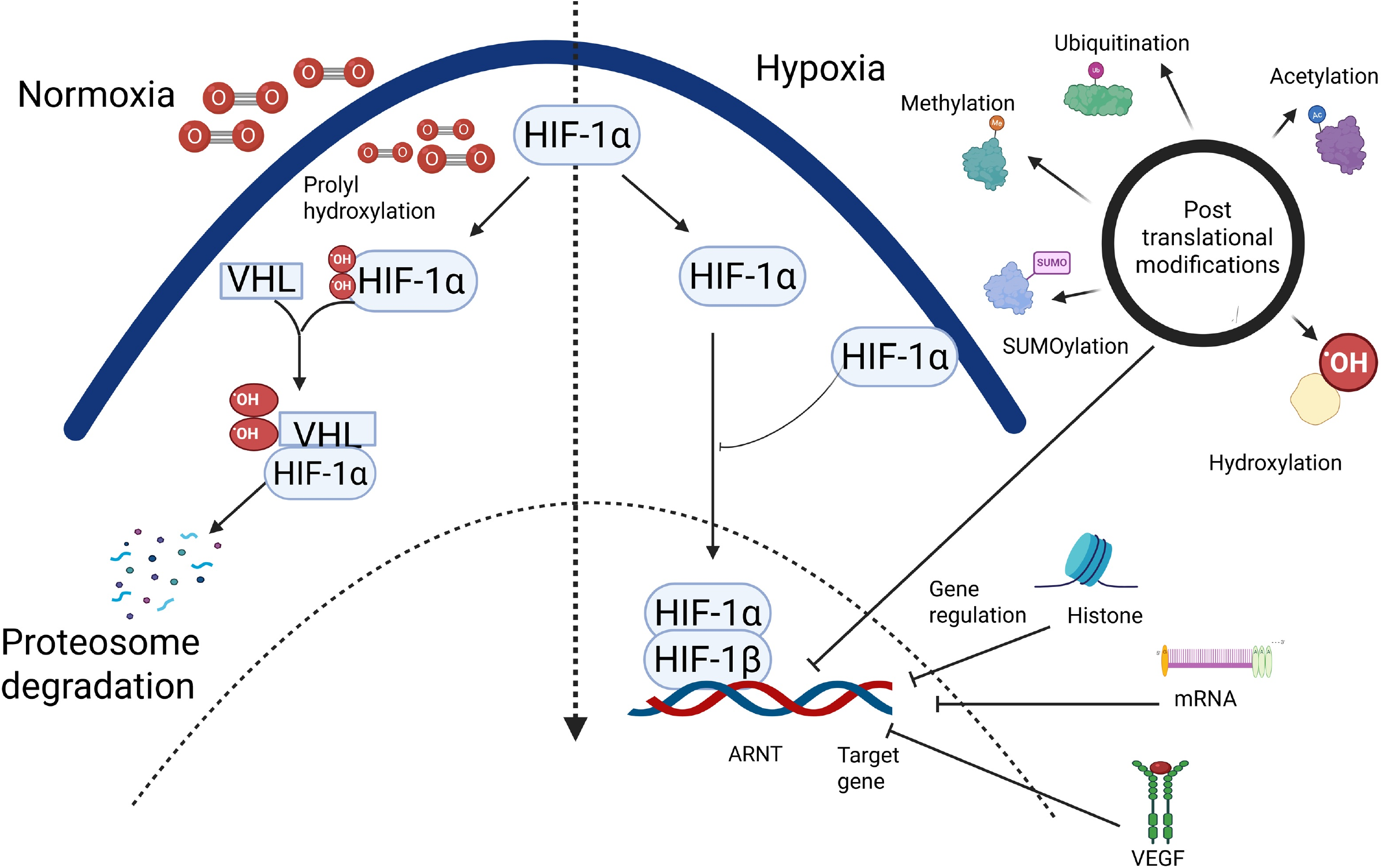
Showing Blocking of HIF pathway by post translational modification leading to reduction in the HIF and HIF induced transcription

Despite high mRNA expression, translation efficiency may be lowered due to methylation induced alteration in the binding of ribosome or regulatory protein in low protein levels. Similarly, histone changes such as acetylation and deacetylation affect chromatin accessibility rather than protein stability. Histone acetylation relaxes chromatin allowing more transcriptional activity whereas deacetylation compact it limiting gene expression. Although these alterations have an impact on transcription, they do not guarantee proportionate protein production since other regulatory mechanism like as miRNA activity and post translational modification are involved (37, 39).

Mutation in HIF1α or EPAS1(HIF2α) mRNA can greatly affect transcript stability and protein production. Certain mutation can cause generation of unstable mRNA transcript which are quickly destroyed before being translated into functional proteins. This instability could lead to formation of premature stop codons which trigger nonsense mediated decay a cellular surveillance process that remove faulty mRNA, mutation induced splicing mistake can result in aberrant transcript variants some of which are degraded due to incorrect exon-exon junctions (40).

Though this study shows a higher mRNA expression and low protein expression in the patients, due to its specific design it is not possible to extrapolate the exact cause of this mismatch in our patients.

## Conclusions

The study identified no significant changes in the expression levels of HIF1α and HIF2α mRNA between OSCC tissues and adjacent normal mucosa, indicating that their overexpression may not be limited to malignancy but also represents physiological adaptions or stress. HIF1α and HIF2α play a vital role in tumor progression and angiogenesis although other variables like gene mutation, epigenetic alteration and normoxic regulation may also contribute to reduced protein expression of HIF1 α and 2 α. Further research into alternative regulatory system with full proteomic profiling is required to understand the exact mechanism of hypoxia and its effector proteins.

## Abbreviation

(HNSCC): Head and Neck Squamous Cell Carcinoma
(OSCC): Oral Squamous Cell Carcinoma
(HIF1α): Hypoxia Inducible Factor 1 Alpha
(HIF1α): Hypoxia Inducible Factor 2 Alpha
(VEGF): Vascular Endothelial Growth Factor
(GAPDH): Glyceraldehyde-3-Phosphate Dehydrogenase
(RT-PCR): Real-Time Polymerase Chain Reaction
(PHD2): Prolyl Hydroxylase 2
(ANG): Angiogenin

## Authors contributions

PS: Conceptualization and original draft preparation, Interpretation of data, Tables, Preparation of manuscript, data collection, literature review, editing of manuscript.

MP: Concept and design, interpretation and editing of manuscript

GN: Concept and design, interpretation of molecular data and presentation of manuscript

DK: Molecular data collection, interpretation and manuscript preparation

AGK: immunohistochemistry data acquisition, interpretation and manuscript perpetration

## Ethical statement

This research was approved by the Ethics Committee, Institutes of Medical Sciences.

## Consent

The written informed consent was taken as per the declaration of Helsinki.

## Acknowledgement

None

## Availability of data

The data is available on reasonable request

